# AAV-DJ Enables Targeted Gene Modulation in Human Ovarian Cells

**DOI:** 10.64898/2026.09.08.749939

**Authors:** Natasha Salpeter, Dilan Gokyer, Bervis Hemis, Amanda K. Wu, Elizabeth L. Tsui, Diane C. Saunders, Meredith Lohman, Anna Kleinhans, Joan K. Riley, Monica M. Laronda, Mazhar Adli, Viviana Gradinaru, Máté Borsos, Elnur Babayev

## Abstract

**Study question:** Can adeno-associated viruses (AAVs) efficiently and safely transduce human primary granulosa cells and human ovarian stromal fibroblasts to modulate the expression of genes involved in follicle recruitment and ovarian microenvironment regulation?

**Summary answer:** AAVs can efficiently transduce human primary granulosa cells and human ovarian stromal cells with low toxicity, with the serotype DJ (AAV-DJ) demonstrating superior performance. AAVs enable targeted modulation of ovarian genes such as *AMHR2* in human primary granulosa cells and *TGF*β*1* in human ovarian stromal cells, supporting the feasibility of gene-based approaches to improve ovarian function.

**What is known already:** Infertility affects approximately 15% of couples, with diminished oocyte quality and quantity being major contributors, particularly with advanced reproductive age. Assisted reproductive technologies rely primarily on hormonal stimulation and do not directly target molecular pathways governing folliculogenesis or the ovarian microenvironment. Animal studies suggest that gene therapy using AAVs offers a promising strategy for tissue-specific and durable gene modulation, yet its application in human ovarian cells remains largely unexplored.

**Study design, size, duration:** This was an experimental laboratory study using primary human granulosa cells from patients undergoing *in vitro* fertilization (IVF) and ovarian stromal cells obtained from post-pubertal patients who donated surplus ovarian tissue from ovarian tissue cryopreservation procedures for research. Sixteen AAV serotypes were evaluated for transduction efficiency, toxicity, and expression dynamics, followed by functional gene knockdown studies using AAV-DJ-based shRNA vectors.

**Participants/materials, setting, methods:** Primary human granulosa cells and ovarian stromal cells were isolated from follicular aspirates and ovarian tissue, respectively. Cells were transduced with GFP-expressing AAV serotypes at varying multiplicities of infection (MOIs), incubation times, and culture days to assess transduction efficiency and cytotoxicity. Lead serotypes were further evaluated in dose-response studies. Functional gene modulation was assessed using AAV-DJ-mediated shRNA knockdown of *AMHR2* in granulosa cells and *TGF*β*1* in stromal cells, quantified by RT-qPCR, immunofluorescence, and western blotting.

**Main results and the role of chance:** Six of sixteen AAV serotypes (AAV2, AAV3, AAV6, AAV9, AAV-DJ, and AAV-X1.1) achieved greater than 10% transduction efficiency in human primary granulosa cells. Among these, AAV-DJ demonstrated the highest efficiency across all MOIs tested, reaching up to 48% transduction at 100,000 MOI, significantly outperforming AAV2 and AAV9. Toxicity increased with viral dose for all serotypes but remained comparable across vectors, with AAV-DJ exhibiting higher efficiency with comparable toxicity. The AAV-DJ-shRNA-*AMHR2* construct produced a significant reduction in expression of *AMHR2* mRNA by ∼81% (p<0.05) in human primary granulosa cells as well as at the protein level in HEK293T-AMHR2-ZsGreen engineered cells (∼85% knockdown, p<0.01). In human ovarian stromal cells, AAV-DJ preferentially transduced EMILIN1 and ACTA2-positive fibroblast and myofibroblast-like cells and enabled robust knockdown of *TGF*β*1* at both transcript (∼89% knockdown, p<0.05) and protein levels (∼67% knockdown, p<0.0001) using AAV-DJ-shRNA-*TGF*β*1* construct.

**Large scale data:** Not applicable.

**Limitations, reasons for caution:** These experiments were performed in vitro using primary human ovarian cells, which may not fully recapitulate the in vivo ovarian environment. Functional reproductive outcomes were not assessed, and future studies will be required to evaluate the effects of gene modulation on folliculogenesis and fertility.

**Wider implications of the findings:** These findings establish AAV-DJ as a promising vector for targeted gene modulation in human ovarian cells. This approach introduces a novel framework for addressing infertility and ovarian function by directly targeting molecular pathways involved in follicle recruitment and ovarian stromal signaling, potentially enhancing current assisted reproductive technologies.

**Study funding/competing interest(s):** This project was supported by Friends of Prentice organization AWD00000757 (E.B.; D.G.), Merkin Translational Research Grant 2023 (M.B. and V.G.). M.M.L is a Gesualdo Family Foundation Research Scholar. E.B. is supported by Women’s Reproductive Health Research (WRHR) Career Development Program K12 HD050121. The authors have declared that no conflict of interest exists.

## What does this mean for patients?

This study looks at whether genes inside human ovarian cells can be safely and precisely modulated as a potential way to improve infertility treatment.

As women get older, fertility declines because the number and quality of eggs decrease and the ovarian environment changes. Current fertility treatments mainly use hormones to stimulate the ovaries, but they do not directly address the biological changes inside ovarian tissue that contribute to age-related infertility.

In this study, ovarian cells were collected from people undergoing fertility treatment or from donated ovarian tissue. We tested whether a virus commonly used in approved gene therapies could deliver instructions into these cells to reduce the activity of specific genes involved in egg activation and age-related changes in the ovary. One viral type was particularly effective at entering key ovarian cell types, and gene activity could be reduced without causing significant cell death.

These findings show that targeted gene modulation in human ovarian cells is possible in the laboratory. This work provides a foundation for future fertility treatments that expand infertility treatment toolbox and move beyond hormone stimulation by directly targeting biological changes in the ovary.

## Introduction

Approximately 15% of couples are impacted by infertility (Ding and Schimenti, 2021, Jiao et al., 2021, Dougherty et al., 2023, Ray et al., 2012, Liu et al., 2021). Most infertility cases can be attributed to a decrease in oocyte quality and quantity as women reach advanced reproductive ages (Chandra et al., 2013). State-of-the-art treatment options for infertility are referred to as assisted reproductive technologies (ART), which show an overall live birth rates per cycle at ∼35% ((CDC), 2024). However, this powerful technology does not overcome the adverse impacts of advanced female age, with live birth rates falling to 15% over the age of 40 and to ∼5% for women over the age of 42 ((SART), 2023). Interestingly, approximately half of infertility cases are attributed to genetic variations (Zorrilla and Yatsenko, 2013); however, no existing treatment targets ovarian gene expression.

The ovary is a highly integrated, multicellular organ in which the oocyte fate is determined by follicle somatic, granulosa and theca cells, as well as by the surrounding stromal compartment (Richards and Pangas, 2010). Ovarian stromal cells, including fibroblasts, myofibroblasts, and vascular cells, regulate extracellular matrix composition, growth factor gradients, tissue stiffness, and angiogenesis, all of which shape the follicular niche (Vong and Kalluri, 2011). Molecular pathways such as the Anti-Müllerian hormone (*AMH)/AMHR2* and transforming growth factor-β (*TGF-*β) signaling play key roles in coordinating follicle recruitment and stromal remodeling (Gowkielewicz et al., 2024, Patton et al., 2021). Dysregulation of these pathways contributes to impaired folliculogenesis, fibrosis, and ovarian aging, suggesting that effective therapeutic strategies must address both granulosa cell signaling and the stromal microenvironment (Fiorentino et al., 2023).

Adeno-associated viruses (AAVs) offer a promising platform for ovarian gene therapy. AAVs are non-pathogenic, have favorable genotoxicity profiles, and support long-term gene expression in human cells (McCarty, 2008). AAV capsids confer serotype-specific tropism, enabling viral vectors to preferentially target distinct cellular populations (Van Vliet et al., 2009). AAV-based therapies are now clinically approved for multiple human diseases, with over 300 ongoing clinical trials and thousands of patients treated for metabolic, neurological, musculoskeletal, and ophthalmologic disorders (Bulcha et al., 2021, Ramamurthy et al., 2022). Although no AAV-based therapies have yet been tested in human reproductive tissues, animal studies provide compelling proof of principle. AAV-mediated gene delivery has been used to restore fertility in luteinizing hormone receptor-deficient male mice (Zhang et al., 2024), and AAV9-mediated delivery of KIT ligand to granulosa cells rescued fertility in infertile female mice (Kanatsu-Shinohara et al., 2022). Together, these studies suggest that AAVs may represent powerful tools for modulating ovarian gene expression in the context of infertility.

Despite these advances, the behavior of AAV vectors in human ovarian cell types remains uncharacterized. In particular, the efficiency, toxicity, and cell-type specificity of different AAV serotypes in human granulosa and stromal cells are unknown, limiting the translation of gene therapy approaches into reproductive medicine. In this study, we systematically evaluated 16 AAV serotypes in primary human ovarian cells to identify vectors capable of efficient and non-toxic gene delivery across key ovarian compartments. We identified AAV-DJ as a leading candidate with superior transduction efficiency and minimal toxicity in human granulosa cells and used this vector to mediate shRNA-based knockdown of *AMHR2*, a central regulator of follicle recruitment (Durlinger et al., 1999, Picard et al., 2017). To extend this strategy to the ovarian microenvironment, we applied the same AAV-based approach to human ovarian stromal cells. Because AAV-DJ preferentially transduced ACTA2 and EMILIN1-positive active stromal subpopulations, we targeted transforming growth factor-β1 (*TGF-*β*1*), a key regulator of stromal remodeling and follicular niche integrity secreted by these cells, to validate functional gene modulation in this compartment (Patton et al., 2021). These studies provide proof-of-concept that coordinated modulation of granulosa and stromal signaling using AAV-DJ-mediated gene delivery may enhance infertility treatment outcomes in future.

## Materials and Methods

### Population

This study was approved by Northwestern University Institutional Review Board (STU00218553). Follicular fluid (FF) was collected between January 2024 and November 2024 from patients undergoing ovarian stimulation and oocyte retrieval for male factor infertility or fertility preservation at the Northwestern Center for Fertility and Reproductive Medicine (NCFRM). Patients were screened for eligibility based on the following criteria: age <35 years, ≥4 follicles measuring >15 mm at the time of retrieval, and no primary female infertility diagnosis. All human ovarian stromal cell samples were obtained as de-identified surplus post-pubertal tissue donated from ovarian tissue cryopreservation procedures for research through Ann & Robert H. Lurie Children’s Hospital of Chicago (Lurie Children’s, IRB #2017-1149 and IRB# 2020-3206).

### Ethical Approval

All participants gave written informed assent and consent according to the protocol approved by Northwestern University Institutional Review Board (STU00218553) and Lurie Children’s Institutional Review Board (IRB #2017-1149 and IRB# 2020-3206).

### Follicular Fluid Processing and Granulosa Cell Cryopreservation

FF was collected from the embryology clinic at NCFRM and immediately underwent processing. Upon collection, FF was centrifuged at 4°C at 400 g for 10 minutes. The pellet was resuspended in 4-5 mL of DMEM/F12 (Cat # 11-320-033), 800 µL of the sample was aliquoted into cryogenic vials (Cat #431386), and 100 µL of DMSO (Cat #219018690) and 100 µL of Fetal Bovine Serum (FBS) (Cat #10082-147) was added. Tubes were sealed and put into a Mr. Frosty Freezing Container (Cat #5100-0050) filled with 100% isopropyl alcohol, then placed into a −80C mechanical freezer for 24 hours which allowed for the controlled cooling of −1C per minute. One day later, cryotubes were transferred to Liquid Nitrogen for long-term storage.

### Thawing, Isolation, Counting and Culturing of Human Primary Granulosa Cells

Human primary granulosa cells were isolated from processed follicular fluid using a density gradient-based method as previously described (Gokyer et al., 2025). Briefly, FBS, L-Glutamine (L-Glut) (Cat #A2916801) and Penicillin Streptomycin (PS) (Cat #15140122) were thawed in a 37°C bead bath. Thaw Media (TM) (DMEM, 10% FBS, 1% PS), Resuspension Media (RM) (DMEM, 5% FBS, 1% PS) and Culture Media (CM) (DMEM, 10% FBS, 4% L-Glut, 1% PS) were prepared in 50 mL conical tubes (Cat #12 565 270). Cryotubes containing processed FF were thawed in a 37°C bead bath. Contents of the cryotubes were transferred to 4 mL of TM, resuspended, and centrifuged at 400 g for 10 minutes at 4°C. The media was aspirated and 4 mL of RM was added to the pellet. Percoll (Cat #P1644-100ML) and PBS (Cat #20012027) were added to the bottom of the conical tube containing RM, at a 8:1:1 to perform density gradient-based granulosa cell isolation. Pipette tip was placed at the very bottom of the tube and Percoll first, then PBS were dispensed slowly. The contents of the tube were centrifuged at 850 g for 10 minutes at 4°C. Centrifuging led to two visible layers with the denser containing many cell types and the less dense, higher up that contained the desired granulosa cells. Granulosa cell layer was carefully removed and placed into a fresh 50 mL conical tube. 5 mL of RM was added onto the isolated cells and the contents were resuspended. 10 µL of Trypan Blue Solution (Cat #T8154100ML) and 10 µL of the cell suspension were combined on a piece of parafilm and 10 µL of this was used for cell counting in a Hemocytometer. The cell suspension was centrifuged at 380 g for 5 minutes at 4°C. In 96-well and 24-well dishes, 30,000 and 100,000 cells were plated, respectively. On Day 1 of culture, the media was aspirated, gently washed with warmed PBS and warmed CM was added. Afterwards, cells were rinsed, and CM was replaced every 2 days, and cultures were used until Day 5 at 37°C.

### Collection and Cryopreservation of Human Ovarian Tissue and Isolation of Interstitial Cells

Ovarian tissue was collected from participants undergoing ovarian tissue cryopreservation for fertility preservation at Lurie Children’s. The ovary was surgically removed via laparoscopic oophorectomy and processed for cryopreservation by thinning the cortical tissue to 1.5 – 2mm thick. The surplus tissue that would not be cryopreserved for patient’s future use was transferred to the lab for research and cryopreserved in CryoStor® CS10 (Cat #07959, STEMCELL Technologies). Cryopreserved tissue fragments were later thawed and digested using a previously published protocol (Wagner et al., 2020, Tsui et al., 2025). Briefly, tissue fragments were thawed for 2-3 minutes in a 37° C water bath, transferred to a 15 mL conical, and alphaMEM + Glutamax (Cat #32-561-037, Fisher Scientific) was added dropwise to dilute CryoStor. Fragments were briefly centrifuged and then transferred to digestion media containing 40 μg/mL Liberase DH (Cat #05401089001, Sigma-Aldrich), 1000 U DNAse I (Cat #10104159001, Millipore Sigma) and 0.4 mg/mL Collagenase IV (Cat #C5138-100mg, Millipore Sigma) in alphaMEM + Glutamax supplemented with 1X insulin-transferrin-selenium (Cat #25-800-CR, Fisher Scientific), 1X antibiotic-antimycotic (Cat #ABL02-100ML, Caisson Labs), and 1 mg/mL human serum albumin (Cat #A9511-5G, Millipore Sigma). Digestion proceeded for a maximum of 45 minutes at 37°C, 5% CO_2_, shaking at 120 rpm, and was quenched by addition of DMEM/F12 supplemented with 10% fetal bovine serum (Cat #F2379-5G, Millipore Sigma) (Tsui et al., 2025). Cells were collected by passing through a 70 µm cell strainer (Cat #229483 CELLTREAT), briefly centrifuged, resuspended in CryoStor® CS10 at approximately 250,000 to 1,000,000 cells per mL, and stored in liquid nitrogen until experimental use.

### Thawing and Culturing of Human Ovarian Stromal Cells

Cryotubes containing isolated stromal cells were thawed in a 37°C bead bath. Cells were resuspended with 7 mL of stromal cell growth media (DMEM/F12, 10% FBS and 1% PS) and centrifuged at 600 xg for 10 minutes at 4°C. Supernatant was aspirated and 4-5 mL of fresh stromal cell growth media was added. Cells were counted using Trypan Blue as described above. In a 96-well plate, cells were cultured at a density of 30,000 cells per well. Stromal cell growth media was replaced the following day and then every 2 days, and cultures were maintained until day 5 at 37°C.

### Plasmid construction and validation

The AAVS1-PRM1_linker_zsGreen1-BSD backbone plasmid (Invitrogen, USA) was used at a stock concentration of 1 mg/mL. The AAVS1-AMHR2_linker_zsGreen1-BSD donor insert was prepared at a concentration of 50 ng/μL. Prior to assembly, plasmid integrity and complete linearization were confirmed by restriction digestion of 2 μg vector DNA using EcoRI-HF (New England Biolabs, USA), performed according to the manufacturer’s instructions. Gibson Assembly was carried out using a 3:1 molar ratio of insert to vector with GeneArt™ Gibson Assembly HiFi Master Mix (Invitrogen, USA). The assembled product was transformed into chemically competent Escherichia coli cells (New England Biolabs, USA), and 50 μL of the transformation reaction was plated onto kanamycin-selective LB agar plates. Following overnight incubation, four single colonies were picked and expanded in selective LB medium for approximately 16 h. Plasmid DNA was isolated using the ZymoPURE Plasmid Miniprep Kit (Zymo Research, USA), glycerol stocks were prepared, and correct construct assembly was confirmed by Sanger sequencing using forward and reverse primers spanning the insert and junction regions.

### Nucleofection and Genome Editing

HEK293T human embryonic kidney cells (ATCC, CRL-3216) (1 × 10) were subjected to nucleofection with 20 μg HiFi SpCas9 protein (Integrated DNA Technologies, USA; Cat. No. 1081061), 4 μg chemically modified sgRNA1, and 4 μg donor plasmid using the SF Cell Line 4D-Nucleofector™ X Kit L (Lonza, Switzerland) and the Lonza Amaxa 4D-Nucleofector system programmed with DS150. Donor plasmids were concentrated by ethanol precipitation and resuspended in P3 buffer prior to nucleofection. After nucleofection, cells were plated in culture medium containing 1 μM AZD-7648 (MedChemExpress, USA), which was replaced with fresh medium lacking the compound after 24 h. Cells were subsequently cultured for 5–7 days before fluorescence-activated cell sorting using a FACSMelody 3-Laser Cell Sorter (BD Biosciences, USA).

### AAV Transduction, Imaging and viability assessment

The 16 AAV capsid serotypes constitutively expressing Green Fluorescent Protein (GFP) upon transduction were generated by the Gradinaru lab at California Institute of Technology, as previously described (Chuapoco et al., 2023). On Day 1 of cell culture 100 µL of warmed Transduction Media (TrM) (DMEM/F12, 5% FBS, 4% L-Glut, 1% PS) was added to cell culture. Using the MOI calculation:, the volume required from the titered AAVs was calculated. At the desired MOI of transduction, the AAVs were added into each well and incubated at 37°C. Each well was imaged using the EVOS Cell Imaging System daily. On Day 3 of culture, the media was gently aspirated and without rinsing, 200 µL of warmed culture media was added. Cultures were kept until Day 5, at which point Hoechst 33342 (Cat#62249) and Propidium Iodide (PI) (Cat #P3566) were added to fresh CM in a 1:1:1000 ratio to perform cell counting and assess cell viability as Hoechst stains all, while PI dead cells only. Cells were incubated at 37°C for 30 minutes and imaged using the EVOS microscope. We used the following formula to quantify the transduction efficiency and toxicity of each AAV serotype at different MOIs.

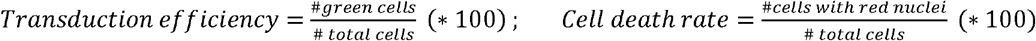

### CellTiter-Glo® 2.0 Cell Viability Assay

The CellTiter-Glo 2.0 Cell Viability Assay (Cat #G9241) was used to measure the cellular ATP content. This assay catalyzes luciferin to oxyluciferin by the luciferase enzyme, which releases energy as luminescence. The luminescence detected is directly proportional to the number of viable cells in the culture. On Day 5 of culture, 200 µL of the CellTiter-Glo assay was added into every well. The plate was mixed on an orbital shaker for 2 minutes, followed by a 10-minute incubation at room temperature. Following incubation, luminescence was measured using a Cytation 3 Plate Reader and Imager at the Analytical bioNanoTechnology Equipment Core (ANTEC) at Northwestern University.

### Immunofluorescent staining of Human Ovarian Stromal Cells

Human ovarian stromal cells were cultured on glass coverslips until day 5 of culture. On day 5, media was aspirated, and coverslips were rinsed with PBS for 2 minutes. Coverslips were then incubated in 4% paraformaldehyde (Cat #157-4-SP) at room temperature for 20 minutes. The coverslips were then washed in PBS for 1 minute and then incubated in 0.5% Triton X-100 (Cat #9036-19-5) at 4°C for 15 minutes. Cells were then washed 3 times for 5 minutes in PBS. The coverslips were blocked in 3% BSA (Cat #A8577) for 1 hour at room temperature. For rabbit host antibodies, 5 mg of goat anti-rabbit IgG (Cat #84300-50497) control was dissolved in 1 mL of H_2_O and further diluted at a 1:100 in 3% BSA. Antibodies: EMILIN1 (Cat #ab197148), FOXL2 (Cat #ab246511) and APOC1 (Cat #ab198288) were prepared at 1:50 ratio in 3% BSA. VWF antibody (Cat #ab154193) was prepared at a 1:250 ratio in 3% BSA. For the mouse host antibodies, 3% BSA was used as a control. Antibodies NR2F2 (Cat #PPH714700) and ACTA2 (Cat #15228-100UL) were prepared at 1:50 ratio in 3% BSA. After blocking, coverslips were incubated with primary antibodies for 1 hour at room temperature. Following this incubation, coverslips were rinsed for 3 times at 5 minutes each in PBS. Goat anti-rabbit IgG, Alexa Fluor 647 (Cat #ab150079) at a 1:2000 ratio in 3% BSA or Donkey anti-mouse IgG, Alexa Fluor 568 (Cat #A10037) at a 1:2000 ratio in 3% BSA were used as secondary antibodies. Coverslips were incubated with the secondary antibody for 1 hour at room temperature in the dark, followed by 3 different 5-minute rinses in PBS. After this final rinse, microscope slides were labelled according to the different well properties, and a drop of DAPI (Cat #H-2000-10) was placed onto each coverslip. The excess PBS was removed gently from each coverslip, and they were placed on the respective microscope slide and sealed with clear quick-dry nail polish. Coverslips were imaged using the TCS SP5 Confocal Microscope using 405 nm, 488 nm and 633 nm excitations.

### Immunoblotting

HEK293T-*AMHR2*-ZsGreen engineered cells were cultured in 6 well plates until Day 4 of culture. Then, media was aspirated, and wells were washed with PBS. Cell plates were frozen in −80°C freezer for 10 minutes to induce cell lysis, and RIPA Buffer Cocktail was prepared at 100:1:1 ratio of RIPA Buffer (Cat #R0278-50ML), Halt™ Phosphatase Inhibitor Single-Use Cocktail (Cat #78420) and Halt™ Protease Inhibitor Cocktail (Cat #87786). 100 µL of RIPA buffer cocktail was added to each well. Cell lysates were collected using a scraper, followed by centrifugation at 16,000 g for 10 minutes at 4°C. The supernatant was transferred to a fresh microcentrifuge tube and stored at 80°C until further use. Protein concentration was determined using a BCA assay kit (Cat #23227). Then, 30 ug protein was diluted with a 1:10 ratio of 2-mercaptoethanol (Cat #M3148) in reducing 4X LDS sample buffer (Cat#NP0007), electrophoresed on a 4% to 12% Novex Bis-Tris polyacrylamide precast gel (Cat #NP0335BOX), and transferred onto polyvinylidene difluoride membrane. The membrane was blocked using 5% non-fat milk (Cat #1706404) for 1 hour at room temperature. Then incubation with ZsGreen monoclonal antibody (Cat #632598) or anti-TGFβ1 primary antibody (Cat #ab215715) was performed at 4°C in 5% non-fat milk overnight. The membranes were then washed and incubated with anti-rabbit IgG HRP linked secondary antibody (Cat #7076S) at a 1:5000 dilution, for 1 hour at room temperature. Detection was performed by West Femto Maximum Sensitivity Substrate (Cat #34096). Then, the membrane was stripped using stripping buffer (Cat #21059), blocked and incubated using HRP conjugated β-Actin Rabbit mAb at a 1:15000 ratio (Cat #5125) in 5% non-fat milk. β-actin was used as a loading control. Immunoblot was quantified using ImageJ software.

### RNA Isolation and Real-Time Quantitative Polymerase Chain Reaction

Total RNA was isolated using the Qiagen RNeasy RNA micro kit (Cat #74004). cDNA was synthesized using qScript cDNA Ultra SuperMix (Cat #76533-174). mRNA levels of *AMHR2*, and *TGF*β*1* were quantified using real-time quantitative polymerase chain reaction and qRT-PCR was normalized to TATA-binding protein (TBP). See supplementary Table 1 for primers.

### Statistical Analysis

The normal distribution of the data was evaluated with the Shapiro–Wilk test. Analysis between the three groups of continuous variables was performed with ordinary one-way ANOVA or Kruskal–Wallis test depending on the distribution. For knockdown experiments, control was set at 1.0 in each replicate and one sample t test was used for comparison. Data are presented as mean ± SEM. *P*-values <0.05 were considered statistically significant. GraphPad Prism version 9.0.1 (Boston, MA, USA) was used for statistical analysis.

## Results

### AAVs exhibit serotype-specific transduction in human primary granulosa cells

We first evaluated whether human primary granulosa cells can be efficiently transduced with AAVs and whether this is serotype dependent. Robust GFP expression was demonstrated in AAV-transduced human primary granulosa cells **(Figure 1A)**. To define optimal transduction conditions, we compared AAV addition, using various serotypes, on day 1 or day 2 of culture (day 0 being the day of plating) with incubation periods of 24 or 48 hours **(Supplementary Figure S1A and B)**. Based on these comparisons, we selected transduction on day 1 of culture with a 48-hour incubation at MOIs of 30,000 and 100,000 for all subsequent screening experiments to minimize experimental variability.

**Figure 1.**
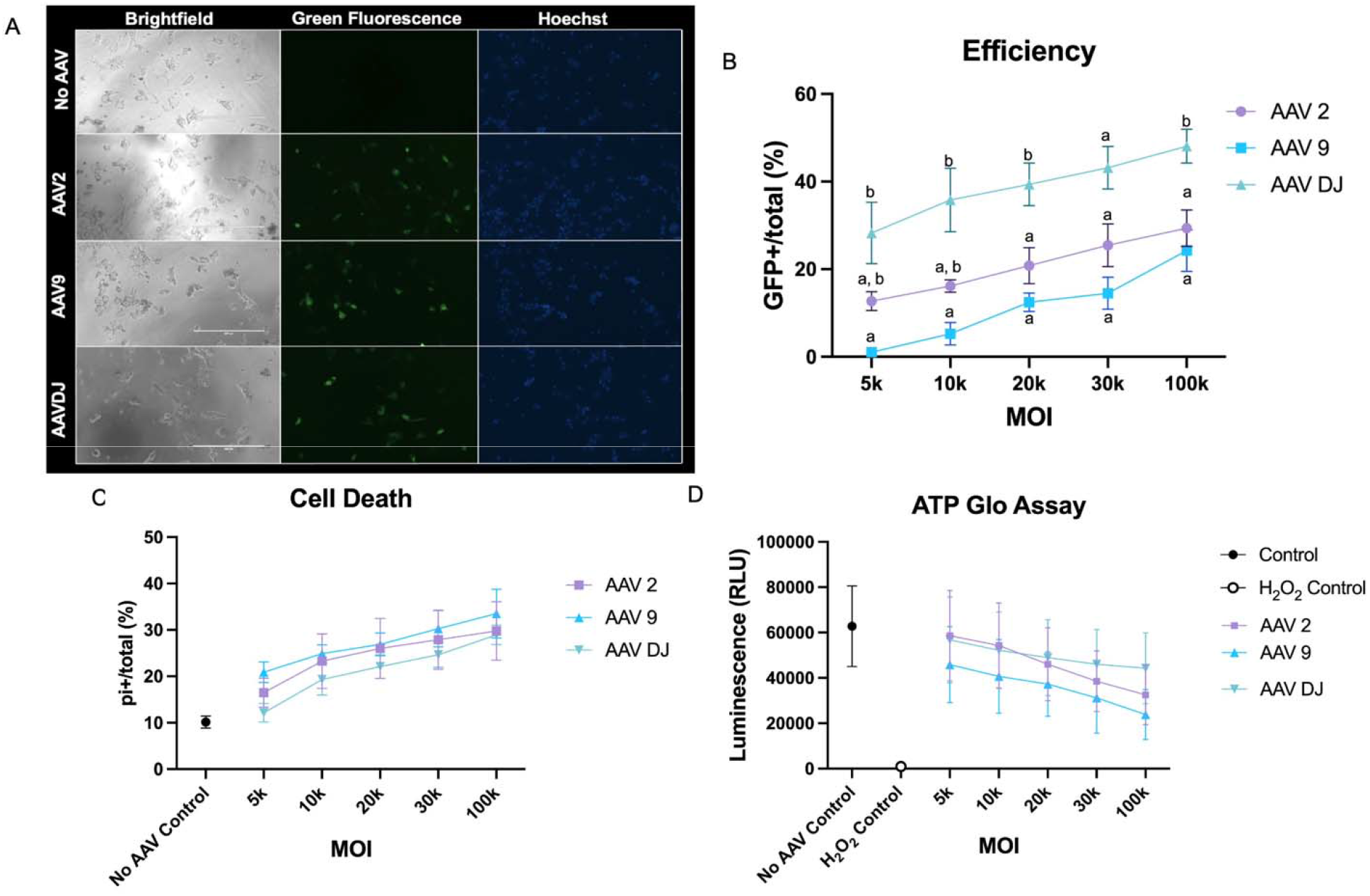
Serotype-dependent AAV transduction in human primary granulosa cells. (A) Representative image demonstrating AAV transduction (green fluorescence due to GFP expression, blue fluorescence, DAPI) in human primary granulosa cells compared to control (contrast and brightness adjusted for visualization and kept the same for all images) (B) Transduction rates on Day 1 of culture at varying MOIs with 48 hours of incubation (dashed line represents 10% transduction efficiency) (C) Intensity trends of GFP signal for Day 1 transduction with 48 hours incubation for the 6 AAV serotypes with >10% cell transduction rates.

Using these conditions, we screened 16 AAV serotypes for their ability to transduce human primary granulosa cells. Transduction efficiency varied across serotypes. We applied a conservative transduction efficiency threshold of greater than 10% to identify candidates for downstream analysis. Six serotypes (AAV2, AAV3, AAV6, AAV9, AAV-DJ, and AAV-X1.1) met this threshold **(Figure 1B)**. For these serotypes, we next examined the dynamics of the GFP expression, tracking the GFP fluorescence intensity in the subsequent days of culture following AAV transduction **(Figure 1C)**. Further screen of these 6 AAVs at low MOIs (5K, 10K and 20K) demonstrated high levels of toxicity (cell death rate >45%) for AAV3 and 6 (**Supplementary Figure S2B**). Additionally, AAVX1.1 which is a modified serotype of AAV9 showed a non-significant trend of lower transduction efficiency compared to AAV9 (**Supplementary Figure S2A**). Therefore, we continued with further testing of top three AAV serotypes (AAV2, 9 and DJ) that demonstrated higher transduction efficiency and lower toxicity.

### AAV-DJ demonstrates the highest transduction efficiency and similar toxicity compared to other serotypes in human primary granulosa cells

To further characterize the transduction efficiency and toxicity of 3 identified AAV serotypes in human primary granulosa cells (**Figure 2A)**, we performed dose response experiments. AAV-DJ showed significantly higher transduction efficiency compared to AAV2 and AAV9 at an MOI of 5000, 10,000, 20,000 and 100,000 (28.26% vs. 12.74% vs. 1.09% p=0.01; 35.81% vs. 20.81% vs. 12.46% p=0.008; 39.39% vs. 20.82% vs. 12.46% p=0.01; 48.05% vs. 29.37% vs. 25.26% p=0.02, respectively) (**Figure 2B**). We also examined AAV cellular toxicity using 2 different assays (PI staining and ATP Glo) (**Figure 2C**, **2D**). All 3 serotypes had similar cellular toxicity across MOIs tested (e.g., 30.29% vs 27.88% vs 24.65% at MOI 30k, p=0.3) with AAV-DJ showing more favorable toxicity trends (**Figure 2D**). Toxicity for all 3 serotypes was similar to controls at 5k MOI but increased with increasing doses (**Figure 2D**). Overall, our experiments indicated AAV-DJ as a promising vector for the granulosa cell gene expression modulation.

**Figure 2.**
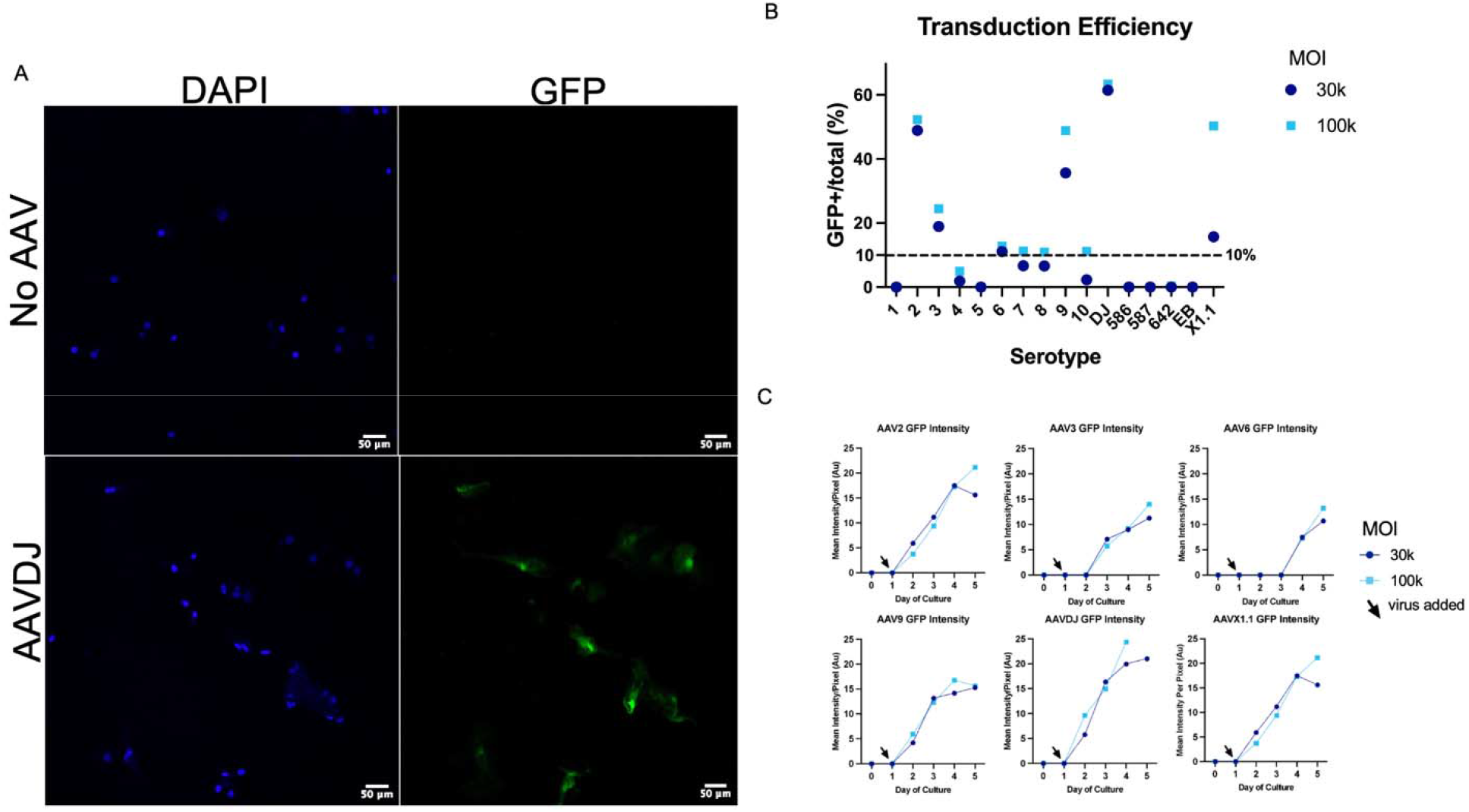
Dose-dependent transduction efficiency and toxicity of AAV2, AAV9, and AAV-DJ in human primary granulosa cells. (A) Representative images of human primary granulosa cells transduced with AAV2,9 and DJ expressing GFP (B) Transduction efficiency dose-response curves (C) Cell death quantified by propidium iodide staining (D) Cellular ATP levels measured by CellTiter-Glo. Values are presented as mean +/- SEM. Points with different letters are significantly different from each other. Cells from three different patients (n=3) were used in each experiment.

### AAV-DJ mediates AMHR2 knockdown in human primary granulosa cells and HEK293T-AMHR2-ZsGreen engineered cell line

To test whether AAV-DJ can modulate the expression of functionally relevant ovarian genes, we targeted *AMHR2*, a key regulator of follicle recruitment. One of the two AAV-DJ-shRNA-*AMHR2* constructs tested led to the significant *AMHR2* knockdown in human primary granulosa cells (∼80% knockdown, p< 0.05) **(Figure 3A)**. As commercial *AMHR2* antibodies proved unreliable, we engineered HEK293T cell line to robustly express *AMHR2* fused to a ZsGreen reporter to enable protein-level knockdown validation. In this system, AAV-DJ-shRNA-*AMHR2* led to a significant decrease in *AMHR2* transcript levels (∼92% knockdown, p<0.0001) **(Figure 3B)** and a corresponding reduction in ZsGreen-tagged *AMHR2* protein detected by anti-zsGreen antibody (∼84% knockdown, p<0.01) **(Figure 3C)**. These data show that AAV-DJ enables *AMHR2* knockdown at both the mRNA and protein levels which can be leveraged to modulate the expression of this gene in human granulosa cells.

**Figure 3.**
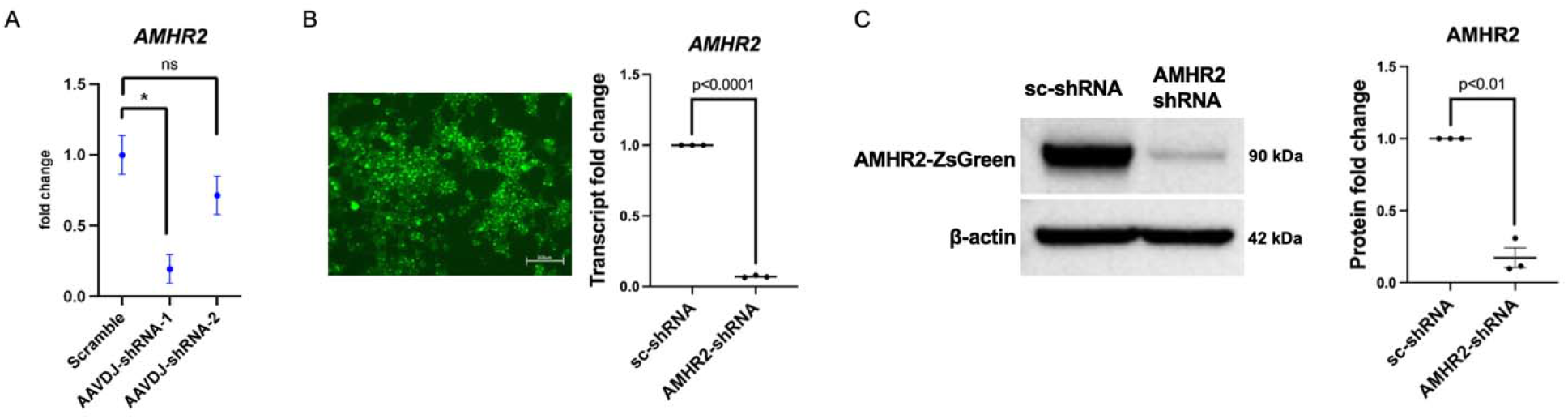
AAV-DJ mediates *AMHR2* knockdown. A) *AMHR2* transcript levels in human primary granulosa cells following transduction with two AAV-DJ-shRNA constructs or scramble control (B) *AMHR2* transcript levels in HEK293T-*AMHR2*-ZsGreen cells following transduction with AAV-DJ-shRNA or scramble control (C) Representative immunoblot (ZsGreen antibody) and quantification for HEK293T-*AMHR2*-ZsGreen cells after AAV-DJ-shRNA knockdown. Cells from three patients (n=3) were used in panel A; bars represent mean ± SEM.

### AAVs can transduce human ovarian stromal cells, fibroblasts and exhibit serotype-specific tropism

To characterize the cell types within the human ovarian stromal cultures, we performed immunofluorescence profiling for key ovarian stromal subtypes. This revealed an enrichment for fibroblasts and myofibroblasts, as indicated by high percentage of cells expressing EMILIN1 (96%), NR2F2 (89%) and ACTA2 (91%), alongside endothelial (VWF) (71%), granulosa (FOXL2) (11%) and theca cells (APOC1) (29%) **(Supplementary Figure S3)**. AAV transduced human stromal ovarian cells demonstrated positive GFP signal **(Figure 4A)**. We screened the six serotypes with the highest transduction efficiency from the granulosa cell analysis, which revealed AAV-DJ exhibiting a trend of stronger GFP signal compared to AAV2, AAV3, AAV6, AAV9, and AAVX1.1 **(Figure 4B)**. Toxicity assays demonstrated overall similar viability between AAV-treated and control cultures **(Figure 4C**, **4D)**. In addition, AAV-DJ-mediated transduction produced a trend of higher GFP signal intensity **(Figure 4E)**. Immunostaining of AAV-DJ transduced human ovarian stromal cells revealed high transduction rate in ACTA2 and EMILIN1 positive cells; among all cells with GFP signal, 75% were ACTA2 and 41% EMILIN1 positive (markers with expected co-expression within the fibroblast lineage), respectively, indicating myofibroblast-like cells as the main AAV-DJ target in human ovarian stroma **(Supplementary Figure S4)**.

**Figure 4.**
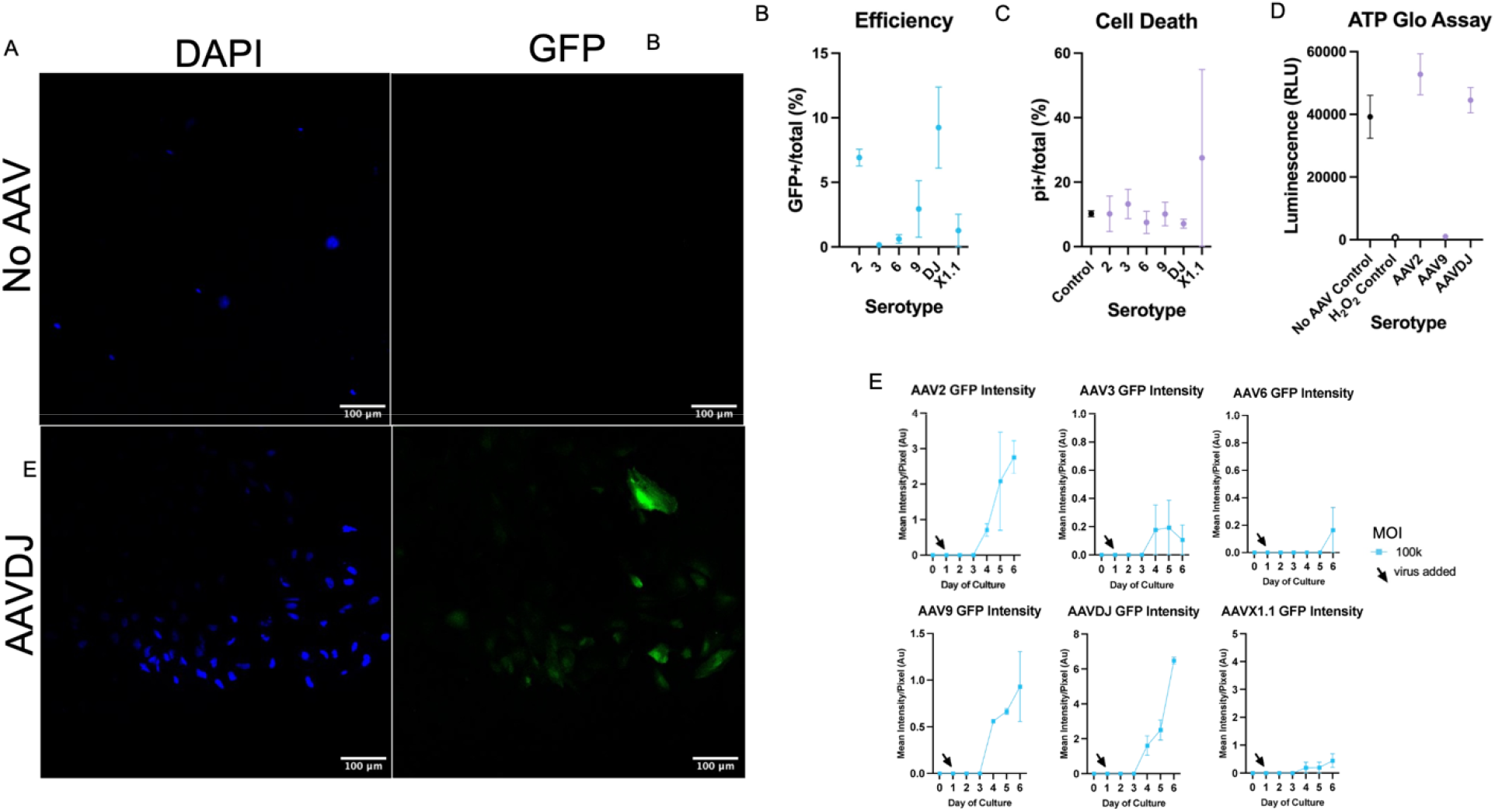
Transduction efficiency of AAV serotypes in human ovarian stromal cells. (A) Representative image demonstrating AAV transduction (green fluorescence due GFP expression, blue fluorescence, DAPI) in human ovarian stromal cells compared to control (contrast and brightness adjusted for visualization and kept the same for all images) (B) Transduction rates and (C, D) cell death in human ovarian stromal cells after AAV transduction (E) GFP signal intensity following AAV transduction in human ovarian stromal cells. Cells from three different individuals (n=3) were used in each experiment. Bars represent mean ± SEM.

### AAV-DJ mediates TGF-β1 knockdown in human ovarian stromal cells

Given the preferential transduction of ovarian myofibroblasts by AAV-DJ, we selected *TGF-*β*1,* a molecule secreted by myofibroblasts which acts as a central regulator of ovarian stromal remodeling and fibrosis (Gu et al., 2024), as a functional target. AAV-DJ-shRNA-*TGF*β*1* led to the significant *TGF*β*1* knockdown at the transcript (∼89% knockdown, p<0.05) (**Figure 5A)**, and the protein level (∼67% knockdown, p<0.0001) **(Figure 5B)**. These data highlight the feasibility of AAV-mediated human ovarian stromal gene expression modulation.

**Figure 5.**
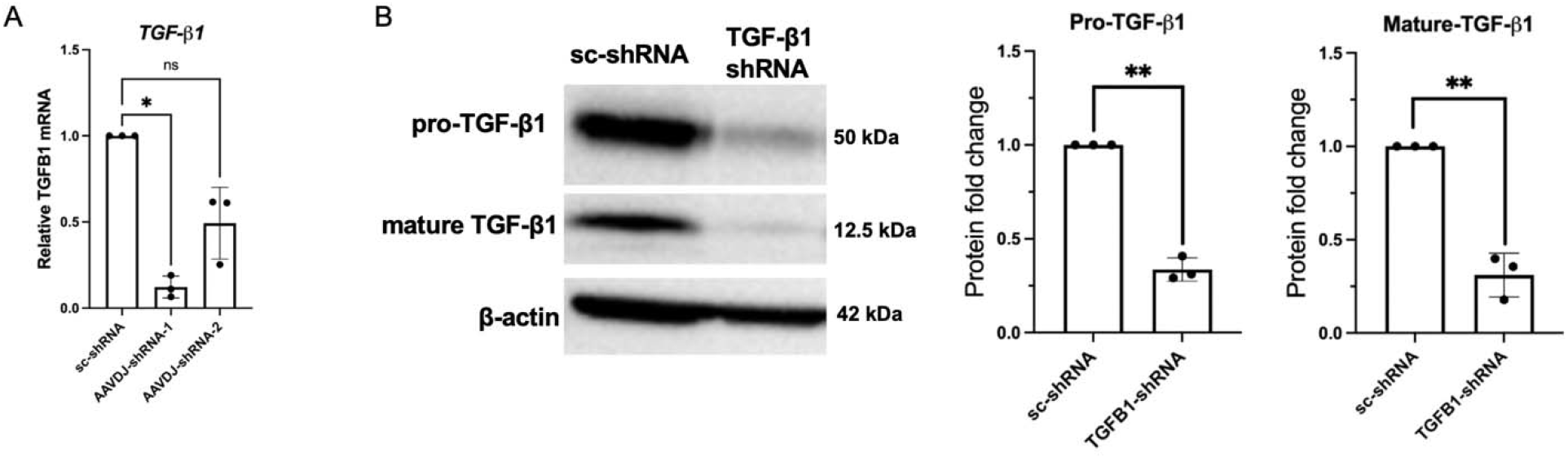
AAV-DJ mediates *TGF*β*1* knockdown in human ovarian stromal cells. A) *TGF*β*1* transcript levels following transduction with two AAV-DJ-shRNA constructs or scramble control in human ovarian stromal cells (B) Representative immunoblots and quantification of precursor and mature *TGF*β*1* protein levels following AAV-DJ-shRNA-knockdown. Cells from three different individuals (n=3) were used in each experiment; bars represent mean ± SD.

## Discussion

ART is a cutting-edge treatment option for women experiencing infertility, yet its efficacy declines significantly for women over the age of 40. Although the process of folliculogenesis is tightly regulated by a complex network of genes, existing infertility treatments do not specifically target these mechanisms (Hsueh et al., 2015). This limitation underscores a critical need for innovative approaches that integrate gene-targeting therapies to improve outcomes for women facing infertility. Moreover, ovarian aging has significant implications for women’s overall health (Liu et al., 2025). Therefore, emergent gene-therapy-based treatments may extend women’s healthspan by prolonging ovarian endocrinological longevity.

AAVs have demonstrated potential in various gene therapy applications. However, their use in human reproductive tissues had not been explored. This study established, for the first time, that human ovarian somatic cells can be efficiently and selectively targeted by AAV-based gene therapy, and that modulation of key ovarian regulatory pathways can be achieved in both granulosa and stromal compartments. By mapping AAV tropism across human ovarian cell types and identifying AAV-DJ as an optimal vector based on its transduction efficiency and toxicity profile, we provide a framework for ovarian gene therapy.

Beyond vector performance, we showed that AAV-DJ supports gene modulation in ovarian cells. In granulosa cells, shRNA-mediated suppression of AMHR2 provides a strategy to modulate AMH signaling, a central regulator of follicle activation and ovarian reserve (Gruijters et al., 2003). In parallel, knockdown of *TGF*β*1* in stromal myofibroblasts demonstrates that this same AAV-DJ-based approach can be extended to the ovarian microenvironment, establishing the versatility of this platform across ovarian cell types.

*AMH* is released by growing follicles, regulates a negative feedback loop that signals to primordial follicles to remain dormant (Carlsson et al., 2006). In fact, *AMH* is being explored as a contraceptive leveraging this mechanism. In mice, AAV9-*AMH* delivered to female mice results in the complete arrest of folliculogenesis (Kano et al., 2017). In cats, overexpression of *AMH* using AAV9-*AMH* constructs provides durable contraception (Vansandt et al., 2023). Conversely, this pathway may be inhibited, allowing increased activation of primordial follicles which has treatment implications for infertility (Durlinger et al., 1999).

In parallel, *TGF*β*1* signalling contributes to age-related ovarian dysfunction through its involvement in stromal fibrosis, extracellular matrix accumulation and restricting the follicular microenvironment (Russ et al., 2020). Increased *TGF*β*1* has been associated with impaired follicle quality and reduced ovarian responsiveness with age (Bertani et al., 2026). Therefore, targeted modulation or silencing of *TGF*β*1* in the ovary may allow for reduced fibrotic remodelling and improved follicle-stroma interactions, both having important implications for infertility treatments to delay or mitigate ovarian aging (He et al., 2025).

By establishing the feasibility of AAV-driven gene expression modulation in human ovarian cells, our studies pave the way for gene therapy–based approaches to treat human infertility and ovarian aging, while also providing a versatile experimental platform for mechanistic studies in human ovarian tissue.

## Authors’ roles

E.B., M.B. and V.G. conceived the original idea and designed the experiments. N.S, M.B., D.G., B.H., D.S., M.L., A.K.W., E.L.T., A.K., J.K.R, carried out the experiments. N.S., M.B., D.G., M.M.L., M.A., V.G., and E.B. analyzed and interpreted the data. N.S. and E.B. wrote the draft. M.B., J.K.R., M.M.L., M.A., V.G., and E.B. provided critical discussion, reviewed, and revised the manuscript.

---

**Supplementary Figure S1.**
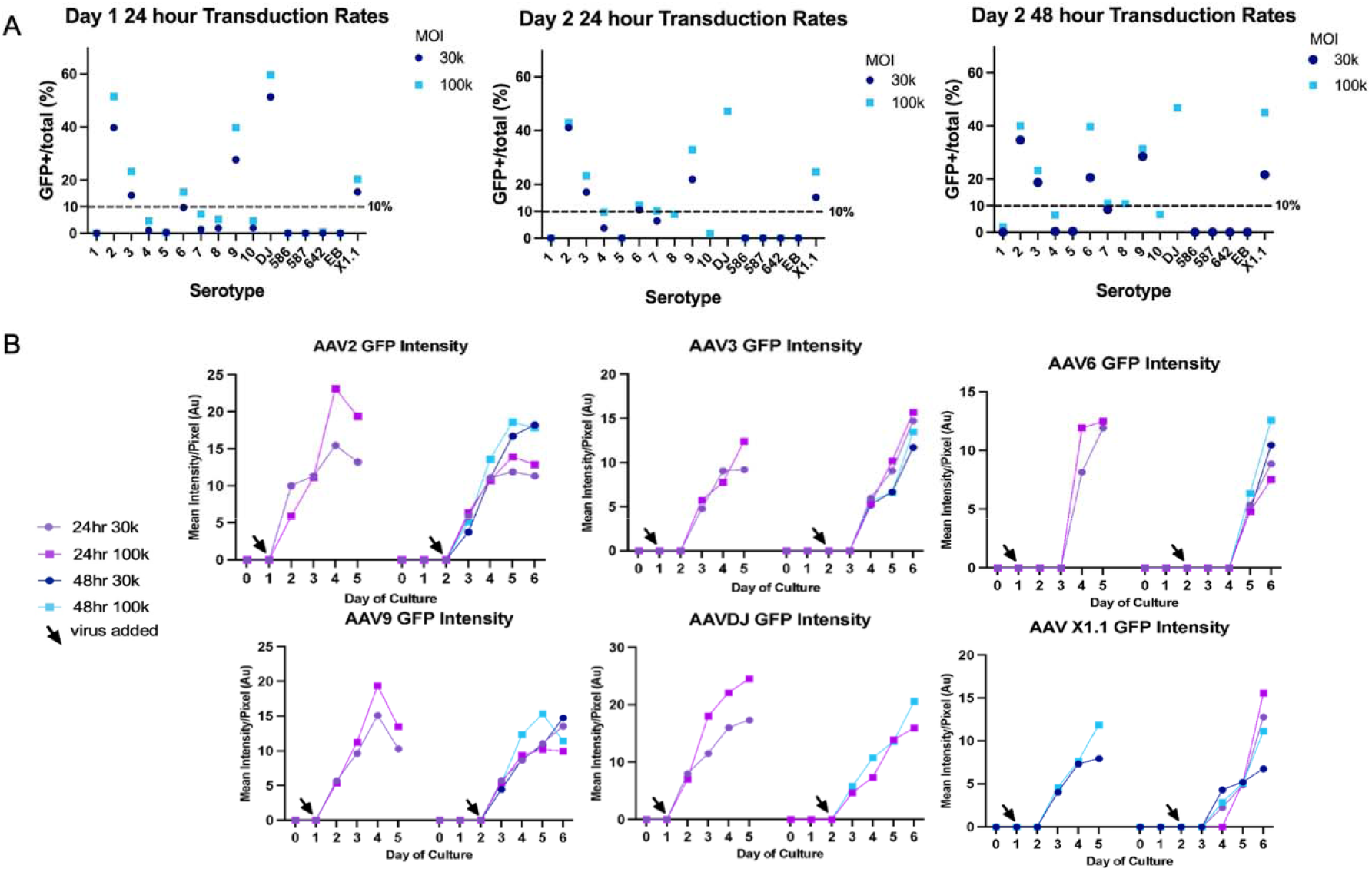
Transduction efficiency of 16 AAV serotypes in human primary granulosa cells. (A) Transduction rates on Day 1 of culture with 24 hours of incubation, Day 2 of culture with 24 hours and 48 hours of incubation at varying MOIs (B) Intensity trends of GFP signal for Day 1 transduction with 24 hours incubation, Day 2 transduction with 24 and 48 hours incubation for the 6 AAV serotypes with >10% cell transduction rates.

**Supplementary Figure S2.**
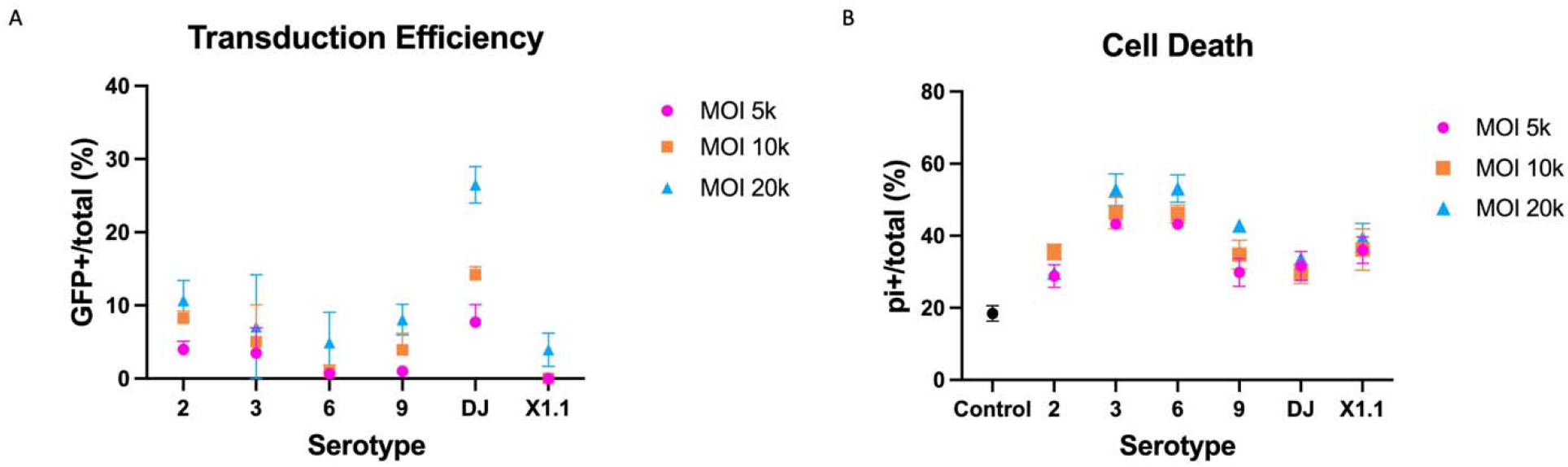
Transduction efficiency and toxicity of 6 AAV serotypes identified in the initial screen in human primary granulosa cells. (A) Transduction rates of AAV 2, 3, 6, 9, X1.1 and DJ at MOIs of 5000, 10,000 and 20,000 (B) Cell death rates of control, AAV2, 3, 6, 9, X1.1 and DJ at MOIs of 5000, 10,000 and 20,000. Cells from three different patients (n=3) were used in each experiment; bars represent mean ± SEM.

**Supplementary Figure S3.**
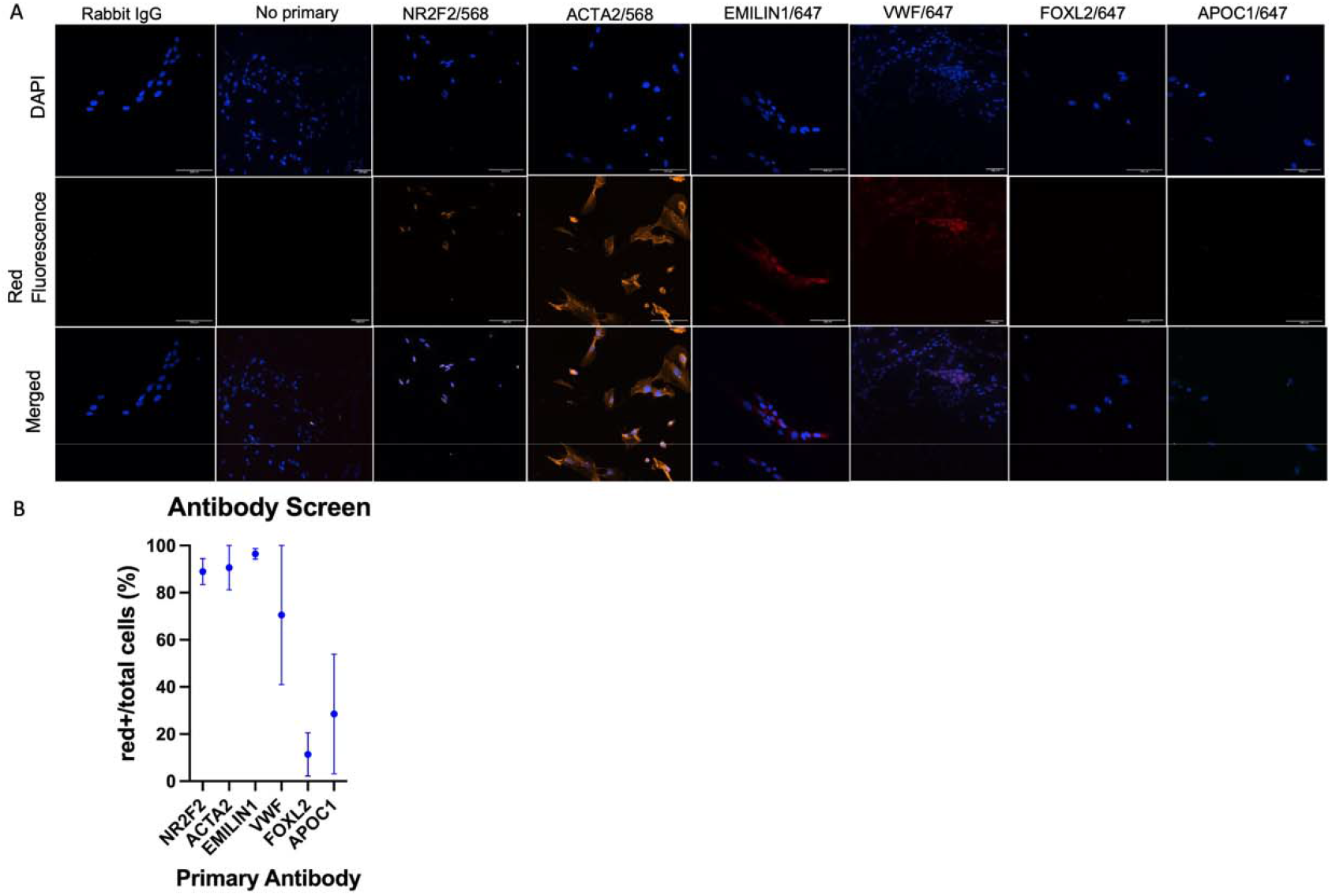
Quantification of different cell types in human ovarian stromal cell cultures. (A) Representative images of NR2F2, ACTA2, EMILIN1, VWF, FOXL2 and APOC1 stained cells (red fluorescence due secondary antibody, blue fluorescence, DAPI) compared to Rabbit IgG and no primary antibody controls (contrast and brightness adjusted for visualization and kept the same for all images) (B) Quantifications of each cell type in culture. Cells from three different individuals (n=3) were used in each experiment. Bars represent mean ± SEM.

**Supplementary Figure S4.**
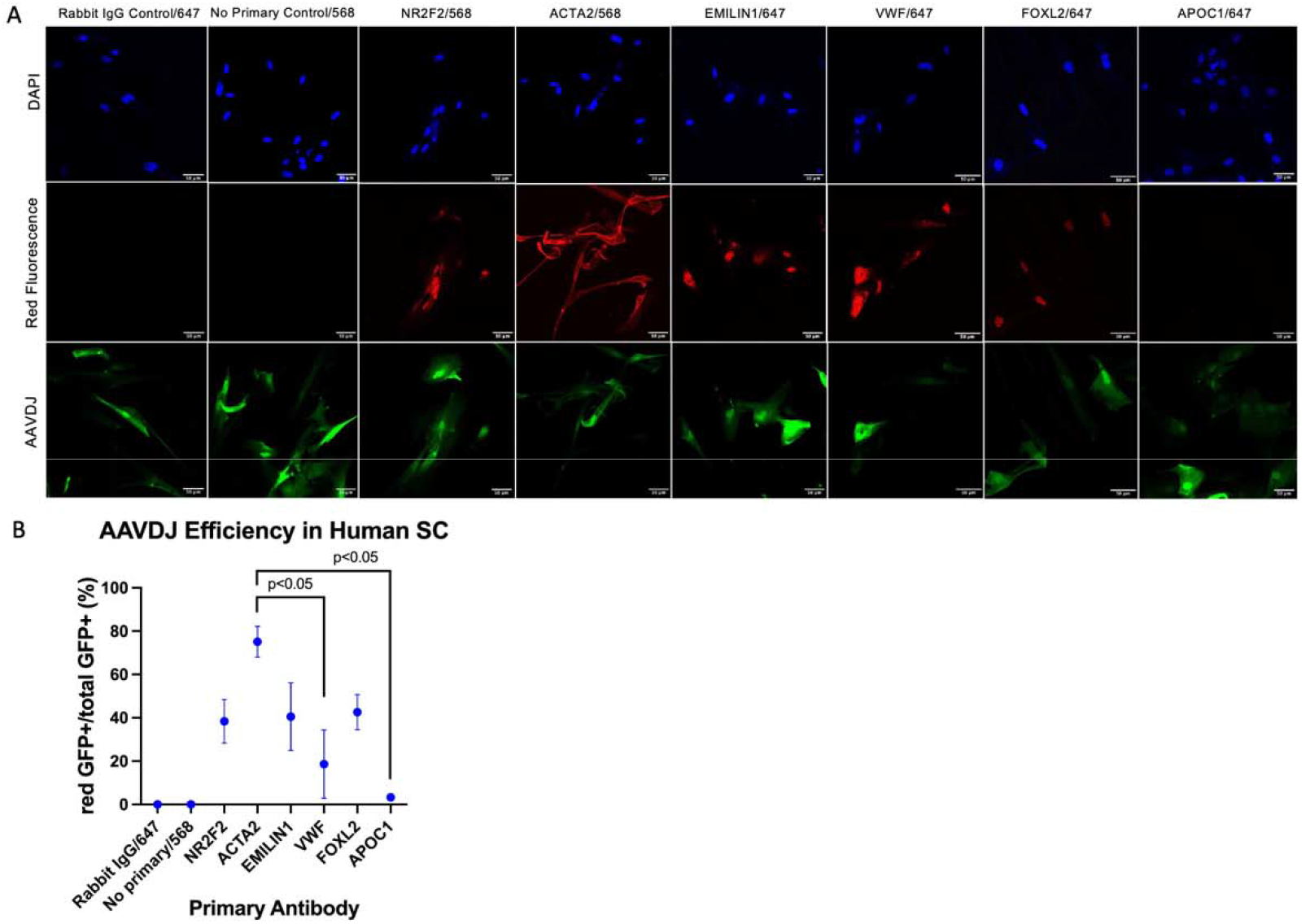
Quantification of AAV-DJ-GFP transduced cell types in human ovarian stromal cell cultures. (A) Representative images of AAV-DJ-GFP transduced cells stained for NR2F2, ACTA2, EMILIN1, VWF, FOXL2 and APOC1 (red fluorescence due secondary antibody, green fluorescence due GFP, blue fluorescence, DAPI) compared to Rabbit IgG and no primary antibody controls (contrast and brightness adjusted for visualization and kept the same for all images) (B) Quantifications of cell types transduced with AAV-DJ. Cells from three different individuals (n=3) were used in each experiment; bars represent mean ± SEM.

**Table S1.**
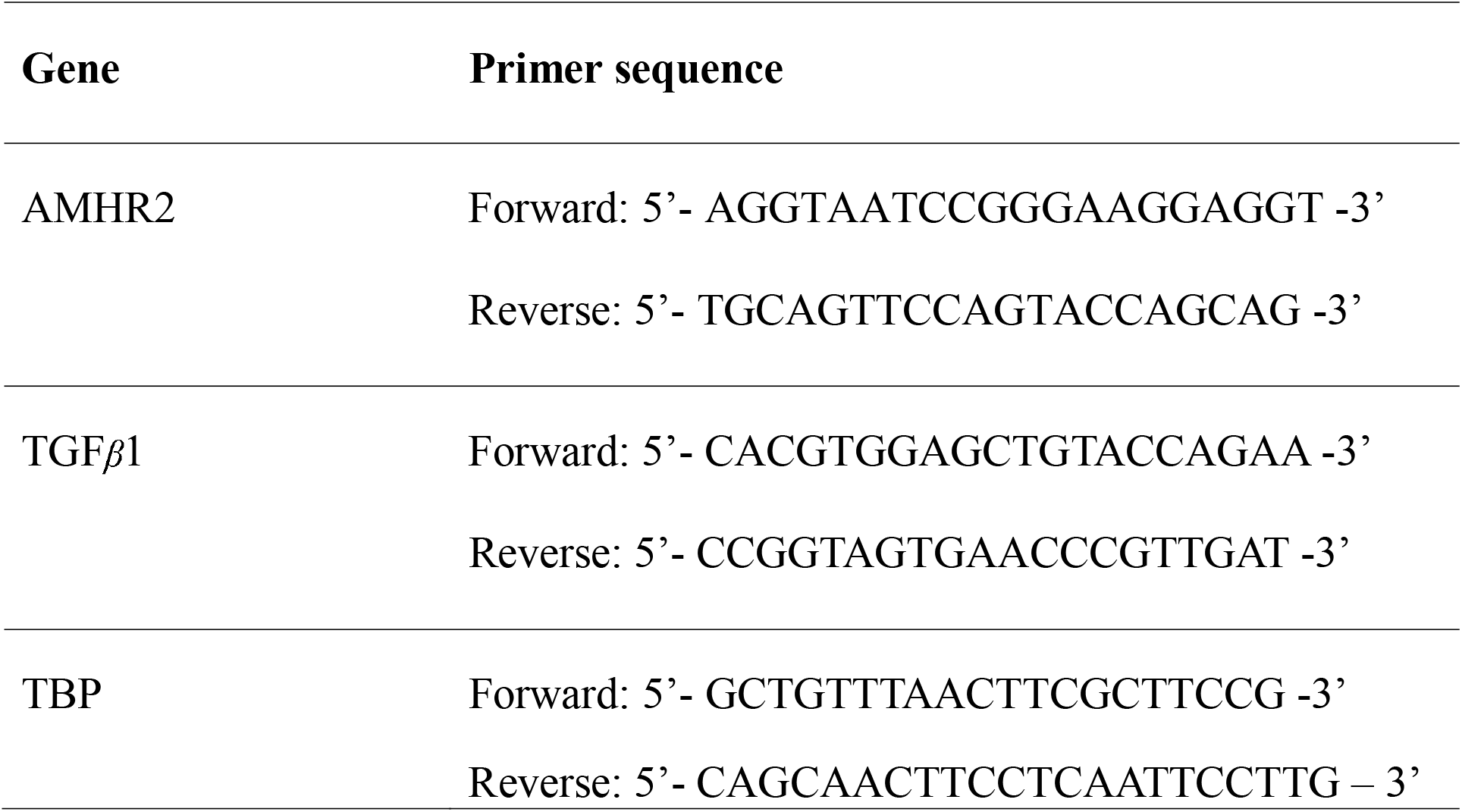
Primer sequences used for qRT-PCR.

**Table S2.** Human Primary Granulosa Cell Donor Information.

| **Anonymous ID** | **Age (years)** | **Diagnosis** | **Serum AMH concentration (ng/mL)** | **Sample type** |
| --- | --- | --- | --- | --- |
| Dn39 | 30.33 | Healthy oocyte donor | 6.09 | Follicular fluid |
| Dn40 | 27.16 | Healthy oocyte donor | 1.99 | Follicular fluid |
| Dn41 | 28.42 | Healthy oocyte donor | 7.83 | Follicular fluid |

**Table S3.** Human Ovarian Stromal Cell Donor Information.

| **Anonymous ID** | **Age**<br>**(years)** | **Diagnosis** | **Previous**<br>**treatment** | **Sample type** |
| --- | --- | --- | --- | --- |
| MLAR-Ov-0213 | 18.34 | Neurofibromatosis (NF1) | No | Tissue fragments from OTC |
| MLAR-Ov-0250 | 19.93 | Hodgkin lymphoma | No | Tissue fragments from OTC |
| MLAR-Ov-0150 | 21.17 | Acute myeloid leukemia | Yes (non-alkylating only) | Tissue fragments from OTC |

## References

(CDC), C. F. D. C. 2024. National Art Summary [Online]. Available: https://www.cdc.gov/art/php/national-summary/index.html [Accessed January 24, 2026].

(SART), S. F. A. R. T. 2023. LIVE BIRTHS PER INTENDED EGG RETRIEVAL (ALL EMBRYO TRANSFERS) [Online]. Available: https://www.sartcorsonline.com/Csr/Public?Clinicpkid=0&reportingYear=2023&newReport=True [Accessed January 24, 2026].

Bertani, N., Alteri, A., Cacciottola, L., D’addato, G., La Sala, G., Lozanoska-Ochser, B., Massimiani, M., Parrella, E., Reggio, A., Russo, E., Campolo, F. & Klinger, F. G. 2026. TGF-beta Signaling in the Pathophysiology of the Ovary: A Double-Edged Regulator. Biomolecules, 16.

Bulcha, J. T., Wang, Y., Ma, H., Tai, P. W. L. & Gao, G. 2021. Viral vector platforms within the gene therapy landscape. Signal Transduct Target Ther, 6, 53.

Carlsson, I. B., Scott, J. E., Visser, J. A., Ritvos, O., Themmen, A. P. & Hovatta, O. 2006. Anti-Mullerian hormone inhibits initiation of growth of human primordial ovarian follicles in vitro. Hum Reprod, 21, 2223–7.

Chandra, A., Copen, C. E. & Stephen, E. H. 2013. Infertility and impaired fecundity in the United States, 1982-2010: data from the National Survey of Family Growth. Natl Health Stat Report, 1–18, 1 p following 19.

Chuapoco, M. R., Flytzanis, N. C., Goeden, N., Christopher Octeau, J., Roxas, K. M., Chan, K. Y., Scherrer, J., Winchester, J., Blackburn, R. J., Campos, L. J., Man, K. N. M., Sun, J., Chen, X., Lefevre, A., Singh, V. P., Arokiaraj, C. M., Shay, T. F., Vendemiatti, J., Jang, M. J., … & Gradinaru, V. 2023. Adeno-associated viral vectors for functional intravenous gene transfer throughout the non-human primate brain. Nat Nanotechnol, 18, 1241–1251.

Ding, X. & Schimenti, J. C. 2021. Strategies to Identify Genetic Variants Causing Infertility. Trends Mol Med, 27, 792–806.

Dougherty, M. P., Poch, A. M., Chorich, L. P., Hawkins, Z. A., Xu, H., Roman, R. A., Liu, H., Brakta, S., Taylor, H. S., Knight, J., Kim, H. G., Diamond, M. P. & Layman, L. C. 2023. Unexplained Female Infertility Associated with Genetic Disease Variants. N Engl J Med, 388, 1055–1056.

Durlinger, A. L., Kramer, P., Karels, B., De Jong, F. H., Uilenbroek, J. T., Grootegoed, J. A. & Themmen, A. P. 1999. Control of primordial follicle recruitment by anti-Mullerian hormone in the mouse ovary. Endocrinology, 140, 5789–96.

Fiorentino, G., Cimadomo, D., Innocenti, F., Soscia, D., Vaiarelli, A., Ubaldi, F. M., Gennarelli, G., Garagna, S., Rienzi, L. & Zuccotti, M. 2023. Biomechanical forces and signals operating in the ovary during folliculogenesis and their dysregulation: implications for fertility. Hum Reprod Update, 29, 1–23.

Gokyer, D., Salpeter, N., Hughes, L., Lee, H. C., Lohman, M. J., Atazhanova, T., Kleinhans, A., Riley, J. K., Pavone, M., Duncan, F. E., Zaniker-Gomez, E. & Babayev, E. 2025. Collection of Human Follicular Fluid, Follicle Somatic Cells, and Immature Oocytes from Individuals Undergoing In Vitro Fertilization. J Vis Exp.

Gowkielewicz, M., Lipka, A., Zdanowski, W., Wasniewski, T., Majewska, M. & Carlberg, C. 2024. Anti-Mullerian hormone: biology and role in endocrinology and cancers. Front Endocrinol (Lausanne), 15, 1468364.

Gruijters, M. J., Visser, J. A., Durlinger, A. L. & Themmen, A. P. 2003. Anti-Mullerian hormone and its role in ovarian function. Mol Cell Endocrinol, 211, 85–90.

Gu, M., Wang, Y. & Yu, Y. 2024. Ovarian fibrosis: molecular mechanisms and potential therapeutic targets. J Ovarian Res, 17, 139.

He, Y., Gan, M., Ma, J., Liang, S., Chen, L., Niu, L., Zhao, Y., Wang, Y., Zhu, L. & Shen, L. 2025. Tgf-beta signaling in the ovary: Emerging roles in development and disease. Int J Biol Macromol, 306, 141455.

Hsueh, A. J., Kawamura, K., Cheng, Y. & Fauser, B. C. 2015. Intraovarian control of early folliculogenesis. Endocr Rev, 36, 1–24.

Jiao, S. Y., Yang, Y. H. & Chen, S. R. 2021. Molecular genetics of infertility: loss-of-function mutations in humans and corresponding knockout/mutated mice. Hum Reprod Update, 27, 154–189.

Kanatsu-Shinohara, M., Lee, J., Miyazaki, T., Morimoto, H. & Shinohara, T. 2022. Adeno-associated-virus-mediated gene delivery to ovaries restores fertility in congenital infertile mice. Cell Rep Med, 3, 100606.

Kano, M., Sosulski, A. E., Zhang, L., Saatcioglu, H. D., Wang, D., Nagykery, N., Sabatini, M. E., Gao, G., Donahoe, P. K. & Pepin, D. 2017. Amh/Mis as a contraceptive that protects the ovarian reserve during chemotherapy. Proc Natl Acad Sci U S A, 114, E1688–E1697.

Liu, J., Tan, Z., He, J., Jin, T., Han, Y., Hu, L. & Huang, S. 2021. Two novel mutations in PADI6 and TLE6 genes cause female infertility due to arrest in embryonic development. J Assist Reprod Genet, 38, 1551–1559.

Liu, X., Zhao, Y., Feng, Y., Wang, S. & Zhang, J. 2025. Ovarian Aging: Mechanisms, Age-Related Disorders, and Therapeutic Interventions. MedComm (2020), 6, e70481.

Mccarty, D. M. 2008. Self-complementary Aav vectors; advances and applications. Mol Ther, 16, 1648–56.

Patton, B. K., Madadi, S. & Pangas, S. A. 2021. Control of ovarian follicle development by TGFbeta family signaling. Curr Opin Endocr Metab Res, 18, 102–110.

Picard, J. Y., Cate, R. L., Racine, C. & Josso, N. 2017. The Persistent Mullerian Duct Syndrome: An Update Based Upon a Personal Experience of 157 Cases. Sex Dev, 11, 109–125.

Ramamurthy, R. M., Atala, A., Porada, C. D. & Almeida-Porada, G. 2022. Organoids and microphysiological systems: Promising models for accelerating Aav gene therapy studies. Front Immunol, 13, 1011143.

Ray, A., Shah, A., Gudi, A. & Homburg, R. 2012. Unexplained infertility: an update and review of practice. Reprod Biomed Online, 24, 591–602.

Richards, J. S. & Pangas, S. A. 2010. The ovary: basic biology and clinical implications. J Clin Invest, 120, 963–72.

Russ, J. E., Haywood, M. E., Mccallie, B. R., Schoolcraft, W. B. & Katz-Jaffe, M. G. 2020. BREAKDOWN OF FOLLICLE-OOCYTE COMMUNICATION CONTRIBUTES TO AGE-RELATED INFERTILITY: DISRUPTED *Tgfb1* SIGNALING IMPAIRS ACQUISITION OF OOCYTE COMPETENCE. Fertility and Sterility, 114, e344.

Tsui, E. L., Saunders, D. C., Lu, L., Kennedy, T., Lee, S., Filip, S. K., Faull, P. A., Hunter, N., Rowell, E. E., Gao, R. & Laronda, M. M. 2025. Primary human ovarian interstitial cells contribute to murine follicle growth through follicle-interstitial paracrine crosstalk. Res Sq.

Van Vliet, K., Mohiuddin, Y., Mcclung, S., Blouin, V., Rolling, F., Moullier, P., Agbandje-Mckenna, M. & Snyder, R. O. 2009. Adeno-associated virus capsid serotype identification: Analytical methods development and application. J Virol Methods, 159, 167–77.

Vansandt, L. M., Meinsohn, M. C., Godin, P., Nagykery, N., Sicher, N., Kano, M., Kashiwagi, A., Chauvin, M., Saatcioglu, H. D., Barnes, J. L., Miller, A. G., Thompson, A. K., Bateman, H. L., Donelan, E. M., Gonzalez, R., Newsom, J., Gao, G., Donahoe, P. K., Wang, D., Swanson, W. F. & Pepin, D. 2023. Durable contraception in the female domestic cat using viral-vectored delivery of a feline anti-Mullerian hormone transgene. Nat Commun, 14, 3140.

Vong, S. & Kalluri, R. 2011. The role of stromal myofibroblast and extracellular matrix in tumor angiogenesis. Genes Cancer, 2, 1139–45.

Wagner, M., Yoshihara, M., Douagi, I., Damdimopoulos, A., Panula, S., Petropoulos, S., Lu, H., Pettersson, K., Palm, K., Katayama, S., Hovatta, O., Kere, J., Lanner, F. & Damdimopoulou, P. 2020. Single-cell analysis of human ovarian cortex identifies distinct cell populations but no oogonial stem cells. Nat Commun, 11, 1147.

Zhang, S., Yang, B., Shen, X., Chen, H., Wang, F., Tan, Z., Ou, W., Yang, C., Liu, C., Peng, H., Luo, P., Peng, L., Lei, Z., Yan, S., Wang, T., Ke, Q., Deng, C., Xiang, A. P. & Xia, K. 2024. Aav-mediated gene therapy restores natural fertility and improves physical function in the Lhcgr-deficient mouse model of Leydig cell failure. Cell Prolif, 57, e13680.

Zorrilla, M. & Yatsenko, A. N. 2013. The Genetics of Infertility: Current Status of the Field. Curr Genet Med Rep, 1.

